# Differential seeding by exogenous R2 and R3 fibrils influences autophagic degradation of intracellular tau aggregates in Tau K18 P301S cells

**DOI:** 10.1101/2023.08.26.554940

**Authors:** Narendran Annadurai, Agáta Kubíčková, Ivo Frydrych, Marián Hajdúch, Viswanath Das

## Abstract

Aggregation of misfolded tau protein is a common feature of tauopathies. Cells employ diverse mechanisms to eliminate misfolded tau with different conformations, contributing to the varied clinical and pathological manifestations of tauopathies. This study focuses on the clearance of seeded tau aggregates following the induction of biosensor cells with R2 and R3 fibrils that exhibit distinct aggregation kinetics and seeding potencies. Our hypothesis is that the dissimilarity in time-dependent intracellular seeding induced by R2 and R3 fibrils underlies the variation in autophagy failure. These discrepancies may account for the heterogeneity of pathology and disease progression between 3R and 4R tauopathies, given the absence of R2 in 3R tau isoforms. In R2-induced cells, alterations in p62 and LC3II/I levels, indicative of proteotoxic stress and autophagy failure, occur sooner than in R3-induced cells. Conversely, LAMP1 levels remained unaffected, suggesting a failure in the fusion of aggregate-containing autophagosomes with lysosomes. This autophagic failure may increase seed-dependent intracellular aggregation in induced cells. Consequently, we assessed the impact of autophagy inducers on the clearance of intracellular tau aggregates in induced cells. Epigallocatechin gallate (EGCG) demonstrated the highest efficacy in inducing autophagy and reducing p62 levels, decreasing seeding and clearing aggregates. Overall, this study elucidates the differential effects of prion-like R2 and R3 strains on autophagy and highlights how compounds like EGCG can selectively reduce tau aggregation to treat specific tauopathies. We provide insights into the distinct mechanisms of autophagy failure and autophagy clearance of intracellular aggregates in cells induced with R2 and R3 fibrils.

## Introduction

Neurofibrillary tangles in tauopathies, including Alzheimer’s disease (AD), are composed of insoluble filaments of the hyperphosphorylated microtubule-associated protein tau [1]. Physiological tau is a highly soluble microtubule-associated protein essential for microtubule assembly and stability [2]. However, insoluble hyperphosphorylated tau aggregates get disseminated in the brain under pathological conditions. Disease severity strongly correlates with the propagation of pathological tau, which is also visible in the clinical phenotypes of tauopathy patients [3,4]. These insoluble aggregates and soluble oligomeric species contribute to neuronal dysfunctions, inflammation and degeneration [5]. Limited proteolysis of pathogenic protein aggregates is critical in the pathogenesis of neurodegenerative diseases [6,7].

While the ubiquitin-proteasome system primarily handles the degradation of soluble and short-lived proteins, autophagy is a dynamic process that breaks down insoluble, long-lived proteins and damaged cellular organelles through lysosomal degradation [8]. Furthermore, autophagy plays a crucial role as a clearance pathway in postmitotic neurons, which, unlike mitotic cells, neurons cannot rely on cell division to alleviate the accumulation of misfolded proteins and damaged organelles [9]. Therefore, autophagy becomes particularly essential in maintaining cellular homeostasis and preventing the buildup of potentially harmful substances in neurons. Defects in the autophagy-lysosome pathway (ALP) are a pathological hallmark of tauopathies [10]. ALP studies using human tauopathy brain samples and animal and cellular models suggest that autophagic vesicles and lysosome accumulation correlate with neuronal toxicity [11]. Neurotoxicity in tauopathies is caused by early tau mislocalisation, oligomerisation, and solubility-related alterations rather than later-stage tau filaments [12,13]. Therefore, early tau deposit clearance may be a promising therapeutic strategy.

Several strategies have been discussed for clearing intracellular tau deposits by small molecules or antibodies disassembling tau aggregates [14]. In P301S tau transgenic mice, primary rat cortical neurons, and N2a cells expressing ΔK280 tau, autophagy induction by trehalose or rapamycin ameliorates disease pathology [15–17]. Moreover, recent studies have revealed that the pathogenic P301L mutation impedes the degradation of tau protein through various autophagy pathways, including macroautophagy and chaperone-mediated autophagy. Conversely, the risk-associated A152T tau mutation guides tau towards distinct autophagy-mediated degradation pathways [18]. These findings underscore the significance of specific autophagy pathways in determining the success or failure of tau clearance under both normal and pathological conditions. Understanding these intricate mechanisms is crucial for unravelling the underlying processes involved in tauopathies and developing potential therapeutic interventions.

Previous reports have indicated that elevated levels of the 4R tau isoform in transgenic mice lead to increased tau hyperphosphorylation and toxicity [19,20]. In chronic traumatic encephalopathy, the early stages of the disease are characterised by higher expression of the 4R tau isoform. However, in later stages with more severe pathology, the 3R tau isoform is expressed at equal or higher levels than the 4R tau isoform [21]. Despite sharing similar clinical features, patients with 4R tauopathies exhibit a shorter disease duration [22] and more aggressive progression [23]. Mutations in 4R tauopathies that result in conformational changes exposing the Repeat 2 (R2) domain, which harbours the VQIINK motif, may contribute to the pronounced pathology associated with 4R tauopathies compared to 3R tauopathies lacking the R2 domain [19,24–26]. We have recently shown that the R2 fibrillizes faster than the Repeat 3 (R3) domain, and R2 aggregates have a greater propensity to seed native tau aggregation in cells than R3 aggregates [27,28]. This increased propensity is unsurprising, as the VQIINK motif in R2 is a more potent driver of tau aggregation than the VQIVYK motif in R3 [24].

Based on these findings, we hypothesised that differences in clearance mechanisms, such as autophagy’s role in eliminating intracellular tau aggregates, may define the progression and pathology of 4R tauopathies. The objective of this study was to investigate how exogenous R2 and R3 fibrils disrupt the ALP and whether activating autophagy could serve as an effective strategy to eliminate intracellular tau aggregates formed after seeding biosensor cells with R2 and R3 aggregates.

## RESULTS

### Exogenous R2 and R3 fibrils affect autophagy and the growth of induced cells

Autophagy induction was measured by quantifying the conversion of LC3-I to LC3-II in fibril-induced cells. A gradual decline in LC3I to -II conversion was observed between 3 to 24 hours post-induction with R2 fibrils (Fig. 1A, C) and up to 48 hours in R3-induced cells (Fig. 1B, D). Cells induced with monomers show LC3I to -II changes similar to uninduced cells, suggesting that monomeric peptides are cleared effectively or do not induce autophagy dysfunctions (Fig. 1A-D). These data indicate that autophagy functions correctly when no or fewer aggregates exist in the cells. When exposed to fibrils, cells become vulnerable to an increased burden of seeding, resulting in autophagy failure to remove the intracellular aggregates. The difference in autophagy timeline between R2 and R3-induced cells might result from the burden of aggregate load, as we have reported that R2 fibrils have a more significant seeding potency than R3 fibrils [28]. The defects in LC3-I/II conversion were also accompanied by increased p62 levels, which showed a trend of time-dependent increase, particularly in R3-induced cells (Fig. 1A, B and E, F). Notably, p62 levels were low in cells induced with monomeric forms of R2 and R3 (Fig. 1A, B). This failure in autophagy in response to the aggregate-induced proteotoxic stress possibly causes the buildup of intracellular aggregates, as observed in our previous study [27,28].

**Figure 1.**
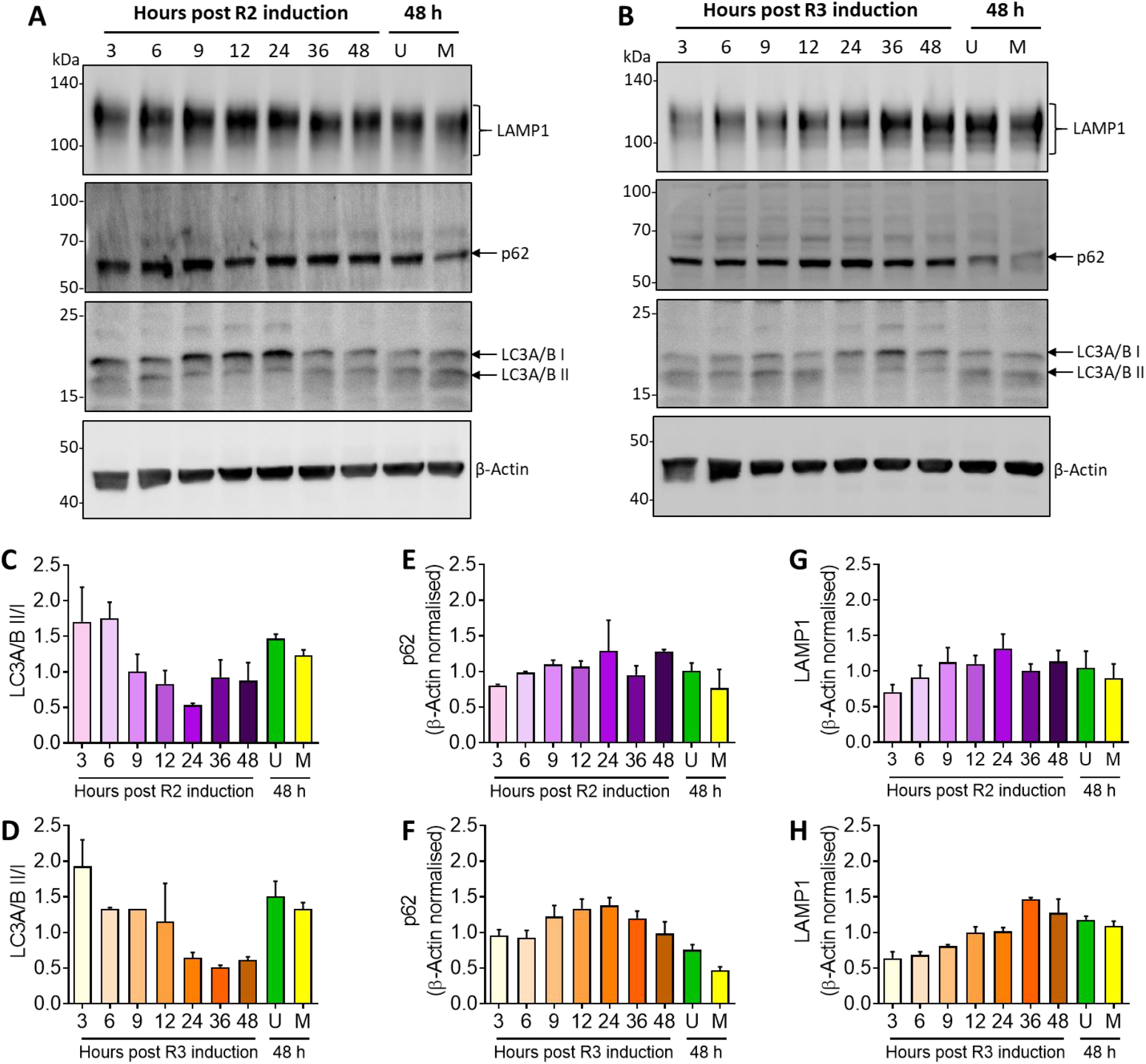
R2 and R3 aggregates effects on autophagy. Western blots (A, B) and quantitative analyses (C-D) show time-dependent changes in LC3A/B I to II conversion (C, D), and p62 (E, F)) and LAMP1 (G, H) levels in biosensor cells expressing P301S Tau following induction with R2 (A, C, E and G) and R3 fibrils (B, D, F and H). Time in hours (h) is the time after the start of transfection with fibrils. U indicates uninduced cells after 48 hours; M indicates cells induced with monomeric forms of R2 and R3 peptides for 48 hours. Data are presented as the mean ± SEM of 2 independent experiments. Images of uncropped blots are shown in the supplemental data (Fig S2).

LAMP1 levels in R2 fibrils-induced cells remained unchanged compared to uninduced cells; however, a time-dependent increasing trend was noted following induction with R3 fibrils (Fig. 1A, B and G, H). These differences between R2 and R3 fibrils-induced cells might also define the distinct seeding efficiency of R2 and R3 fibrils. Overall, the data indicate that autophagosome production is impeded or that the aggregate load is not delivered to lysosomes for degradation in induced cells. Alternatively, unprocessed aggregates may escape from autophagosomes and seed the nearby cells [29]. The overall failure of autophagy processes on the clearance of intracellular P301S tau aggregates in R2 and R3 fibrils-induced cells defines the impact of tau mutations, as P301L mutation was found to impair the autophagy-mediated degradation process[18].

Mitotic cells containing aggresomes of misfolded ubiquitinated proteins can go through mitosis normally, resulting in the asymmetric distribution of aggregates to one of the daughter cells. [30]. We previously reported a reduction in the growth of cells induced with R3 fibrils [31]. Therefore, we examined phospho-Histone H3 (Ser10) levels in the cell population sorted based on 0, 50 and 100% intracellular aggregate positivity (Ag+) after induction with R2 and R3 (Fig. 2A). A phospho-Histone H3 (Ser10) decrease was observed in induced cells with 100% Ag+ (Fig. 2B), suggesting decreased proliferation under proteotoxic stress. The results align with our earlier study, showing reduced growth in cell cultures induced with R3 fibrils [27].

**Figure 2.**
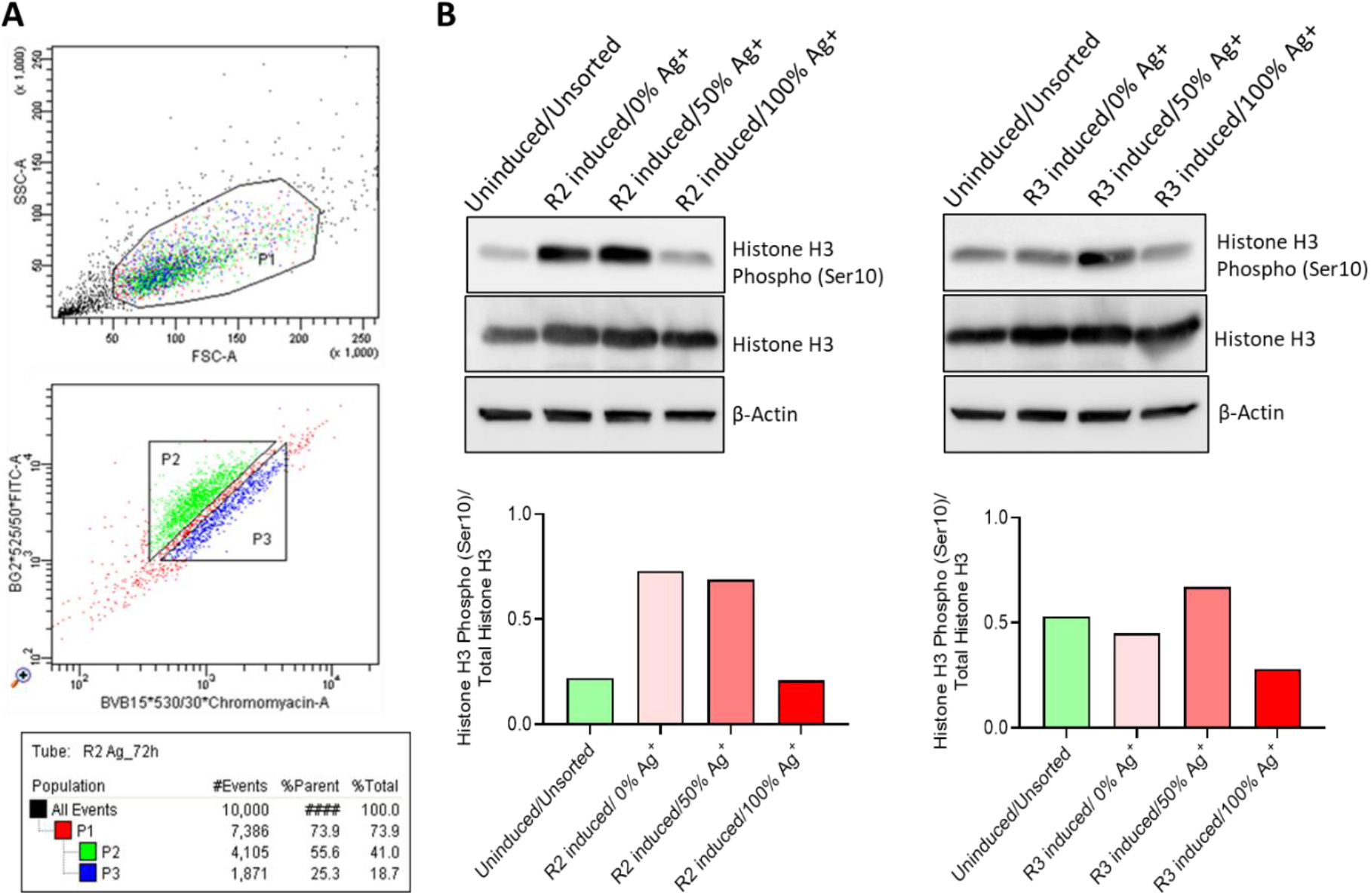
Cell sorting and western blot of mitotic marker, histone h3 phospho (ser10) levels in seeded cells. (A) FACS sorting plots showing intact live cells (P1) and cell population with 0% intracellular Ag+ (P2) and 100% intracellular Ag+ (P3). (B) Changes in Histone H3 Phospho (Ser10) levels in uninduced/unsorted cells and cells induced with 100 nM R2 and R3 for 72 hours and sorted based on 0%, 50% and 100% intracellular Ag+. The fraction containing 50% Ag+ was prepared by mixing 0% and 100% Ag+ collected fractions in equal ratios. Images of uncropped blots are shown in the supplemental data (Fig S3).

### Enhancing autophagy rescues cells from intracellular seeding by R3 and R2 fibrils

Since exogenous fibrils resulted in ALP defects, we next investigated whether pharmacological interventions could rescue cells by clearing intracellular aggregates of seeded tau. We selected drugs that have been reported to alter the ALP through different mechanisms [32] and chose non-cytotoxic concentrations to exclude any potential bias resulting from off-target cytotoxicity (See Fig. S1 for cytotoxicity evaluation of drugs). Cells were induced with fibrils and treated with drugs sequentially or concomitantly. In sequential treatment, cells were first induced with fibrils for 6 h or 48 h, followed by washing to remove residual fibrils and then treated with drugs for 12 h or 72 h. In concomitant treatment, cells were given fibrils and drugs for 24 h.

Epigallocatechin gallate (EGCG) at all tested concentrations significantly reduced seeding by R2 and R3 in both types of treatments (Fig. 3A, B). The effects of resveratrol (RSV) on seeding were more pronounced when administered sequentially following R2 and R3 induction. Additionally, concomitantly treating biosensor cells with R2 and 20 µM RSV significantly reduced seeding (Fig 3A). Among other tested drugs, l-ascorbic acid (LAA) and quercetin (QCT) significantly reduced seeding when administered sequentially after the induction of cells with R3. The autophagy inhibitor chloroquine (CQ) significantly increased seeding under concomitant treatment at the highest tested concentration of 2 µM and showed a non-significant trend of dose-dependently increasing seeding in R3-induced cells (Fig 3A). CQ blocks the fusion of autophagosomes and lysosomes, enhancing the rate of failure of autophagy flux in fibril-induced cells that might explain the increased intracellular seeding compared to untreated cells [33].

**Figure 3.**
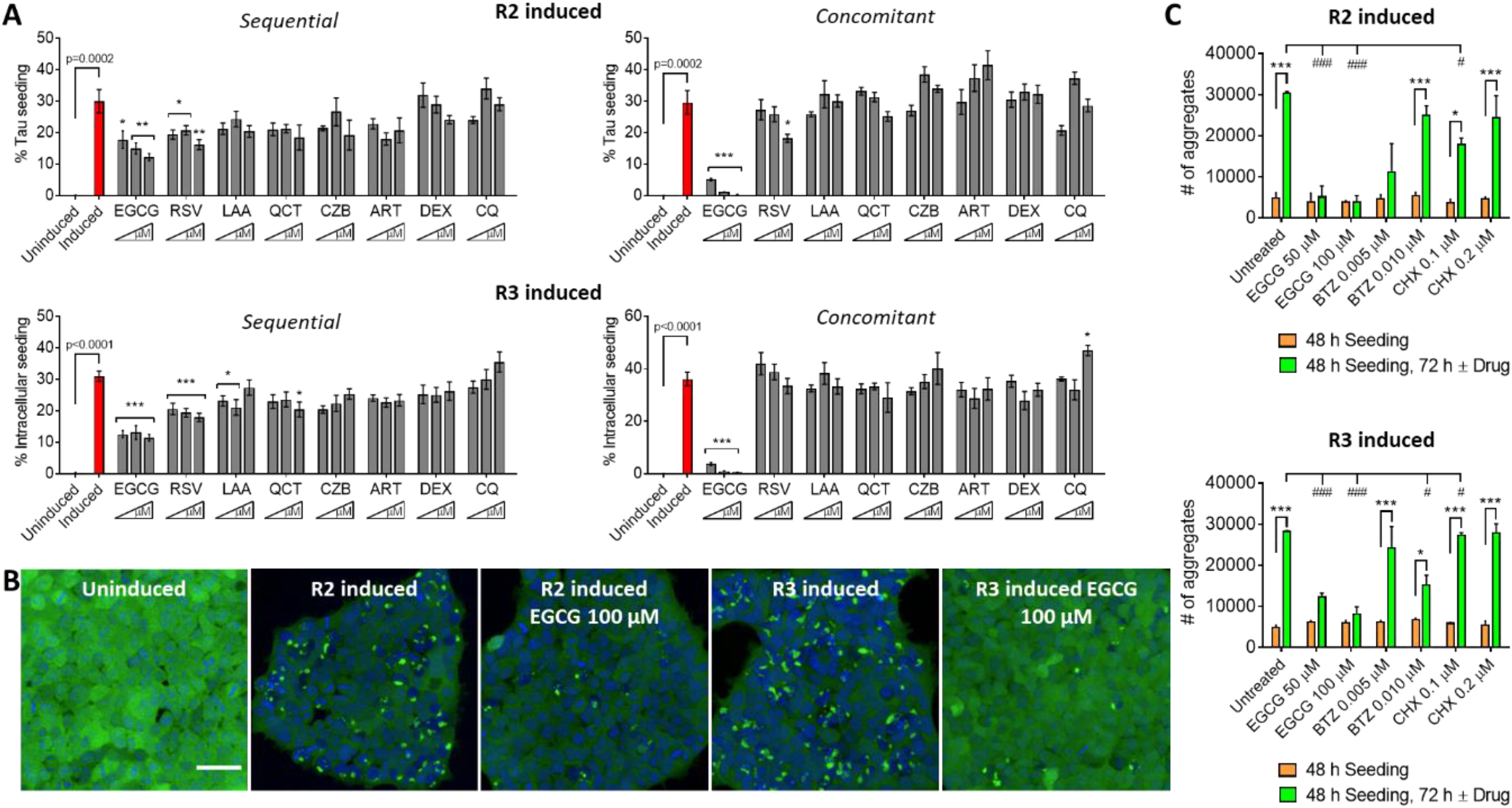
Early autophagy enhancement reduces intracellular tau seeding. (A) Percentage seeding in uninduced cells and cells induced with fibrils without drug treatment and cells after sequential and concomitant treatment with fibrils and autophagy activators/inhibitors. EGCG (25, 50 and 100 µM); RSV (1, 10 and 20 µM); L-Ascorbic acid, LAA (10, 25 and 50 µM); Quercetin, QCT (0.1, 1 and 2 µM); Carbamazepine, CZB (1, 5 and 10 µM); Artemisinin, ART (1, 5 and 10 µM); Dexamethasone, DEX (1, 5 and 10 µM); Chloroquine, CQ (0.1, 1 and 2 µM). Mean ± SEM, n = 2-3. Uninduced vs Induced, Unpaired *t*-test, Two-tailed. ***p<0.001, *p < 0.05, Drug-untreated induced cells (red column) vs Drug-treated induced cells (grey columns), one-way ANOVA, Dunnett’s posthoc test. (B) Uninduced and induced cells with intracellular aggregates of seeded native tau (green, fluorescent inclusions, and EGCG-mediated decrease in seeding after 12 h of treatment. Scale bar: 50 μm. (C) Biosensor cells were induced with R2 or R3 for 48 h, followed by treatment with EGCG, bortezomib (BTZ) and cycloheximide (CHX) for 72 hours. Mean ± SEM, n = 2, two-way ANOVA, Sidak’s multiple comparisons test.

In the above section, we show the effective clearance of intracellular tau aggregates by EGCG when it was added at the beginning or early phase of the seeding (Fig. 3A). Subsequently, we evaluated the efficiency of autophagy induction at a late stage of seeding, when more intracellular aggregates had already accumulated in cells, thus mimicking therapeutically relevant situation. We first induced cells with R2 and R3 fibrils for 48 hours, washed the cells to remove any residual fibrils, and then treated induced cells with EGCG, bortezomib (BTZ) and cycloheximide (CHX), which is known to inhibit autophagy [34] for 72 hours (Fig. 3C). Compared to untreated cells, EGCG reduced intracellular aggregates in both R2 and R3 induced cells (Fig. 3C). However, it was unable to reduce intracellular aggregates below the threshold seeding levels noted after 48 hours of cell induction with fibrils. This suggests that aggregates are more resistant to autophagic clearance at the late phase of the seeding. Early clearance of aggregates before a major aggregate load buildup might help in APL-mediated clearance of misfolded aggregates.

### EGCG effect on ALP proteins in cells induced with R2 and R3 fibrils

To show that exogenous fibril-induced seeding of native tau can be mitigated by enhancing autophagy, we examined ALP-associated proteins after 48 hours of treatment with autophagy inducers. We chose EGCG and RSV for further studies because they had had a better effect in reducing seeding than other tested autophagy inducers (Fig. 3). EGCG dose-dependently increased LC3A/B-I to -II conversion in both R2 and R3-induced cells (Fig. 4A, B), whereas the effect of RSV was evident in R2-induced cells (Fig. 4C, D). We next looked at p62 levels to see if EGCG clearance of intracellular aggregates reduced proteotoxic stress. As noted earlier, R2- and R3-induced cells displayed proteotoxic stress (Fig. 1A). EGCG significantly reduced this stress, as indicated by the decrease in p62 levels (Fig. 4A, B). However, there was no significant change in p62 levels after RSV treatment of cells (Fig. 4C, D). These results confirm that EGCG-mediated autophagy reduces proteotoxic stress, thereby decreasing intracellular tau seeding. Additionally, LAMP1 levels increased after EGCG and RSV treatment in R2 or R3-induced cells, indicating that EGCG mediated the clearance of intracellular aggregates by enhancing autophagy flux. A similar result was noted in primary cortical neuronal cultures, where EGCG cleared pathological tau inclusions by enhancing LAMP1 levels [35]. These data show that EGCG is a potent autophagy inducer that clears intracellular aggregation in cells induced by both R2 and R3 fibrils. Although EGCG was an effective inhibitor of intracellular aggregation, the concentrations needed for biological effects are difficult to achieve in clinical settings, and there is a need to search for more potent compounds.

**Figure 4.**
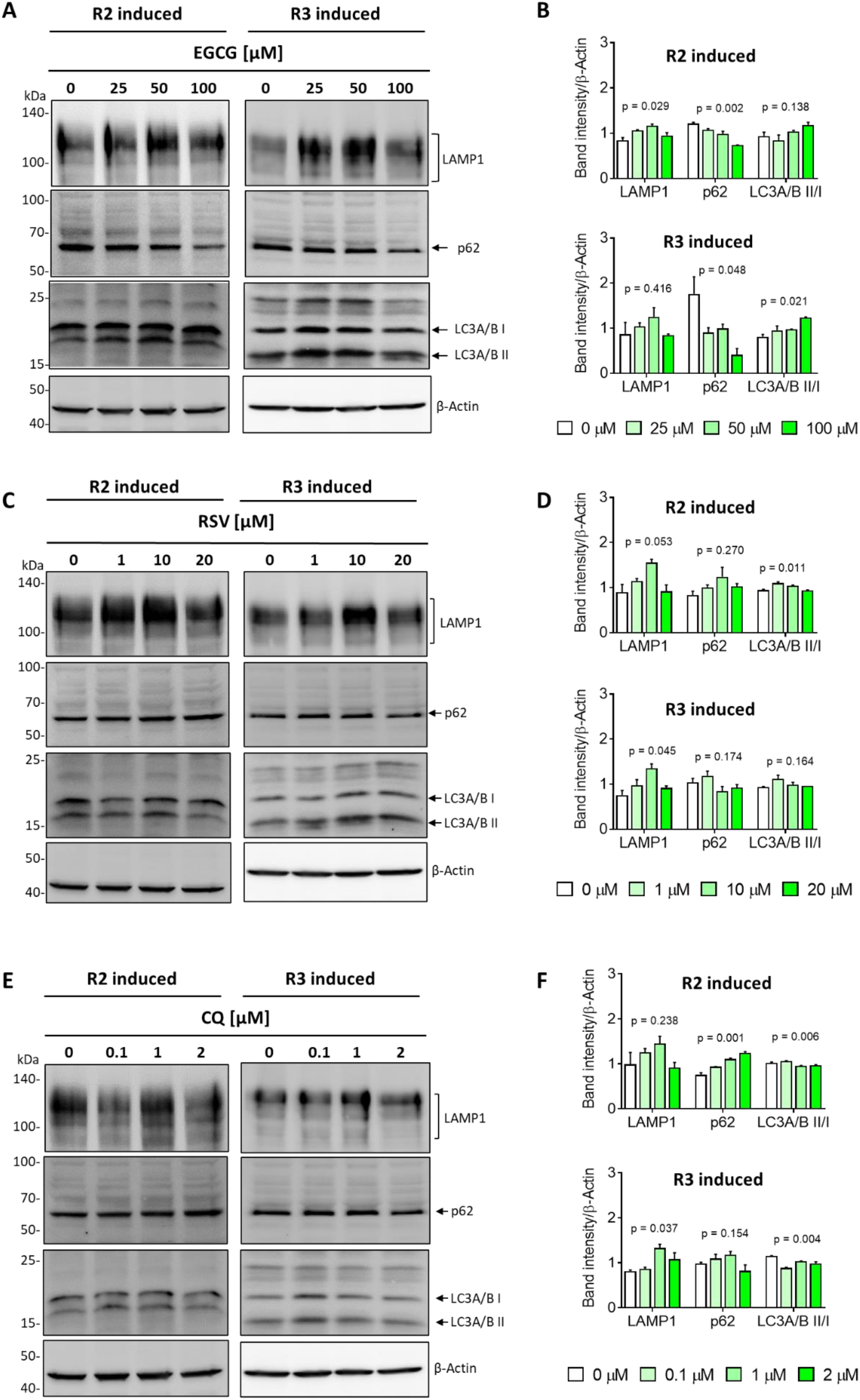
ECGC clears proteotoxic stress in R2 and R3 fibril-induced cells. (A-F) Representative Western blots and quantification showing changes in LAMP and p62 levels and LC3-I to LC3-II conversion in cells induced with R2 or R3 fibrils after 48 h of treatment with three concentrations of EGCG (A, B), RSV (C, D) and CQ (E, F). Mean ± SEM, n = 2 independent experiments. P values summary of one-way ANOVA analysis (0 µM vs drug treatment) are shown. Images of uncropped blots are shown in the supplemental data (Fig S4).

**Figure 5.**
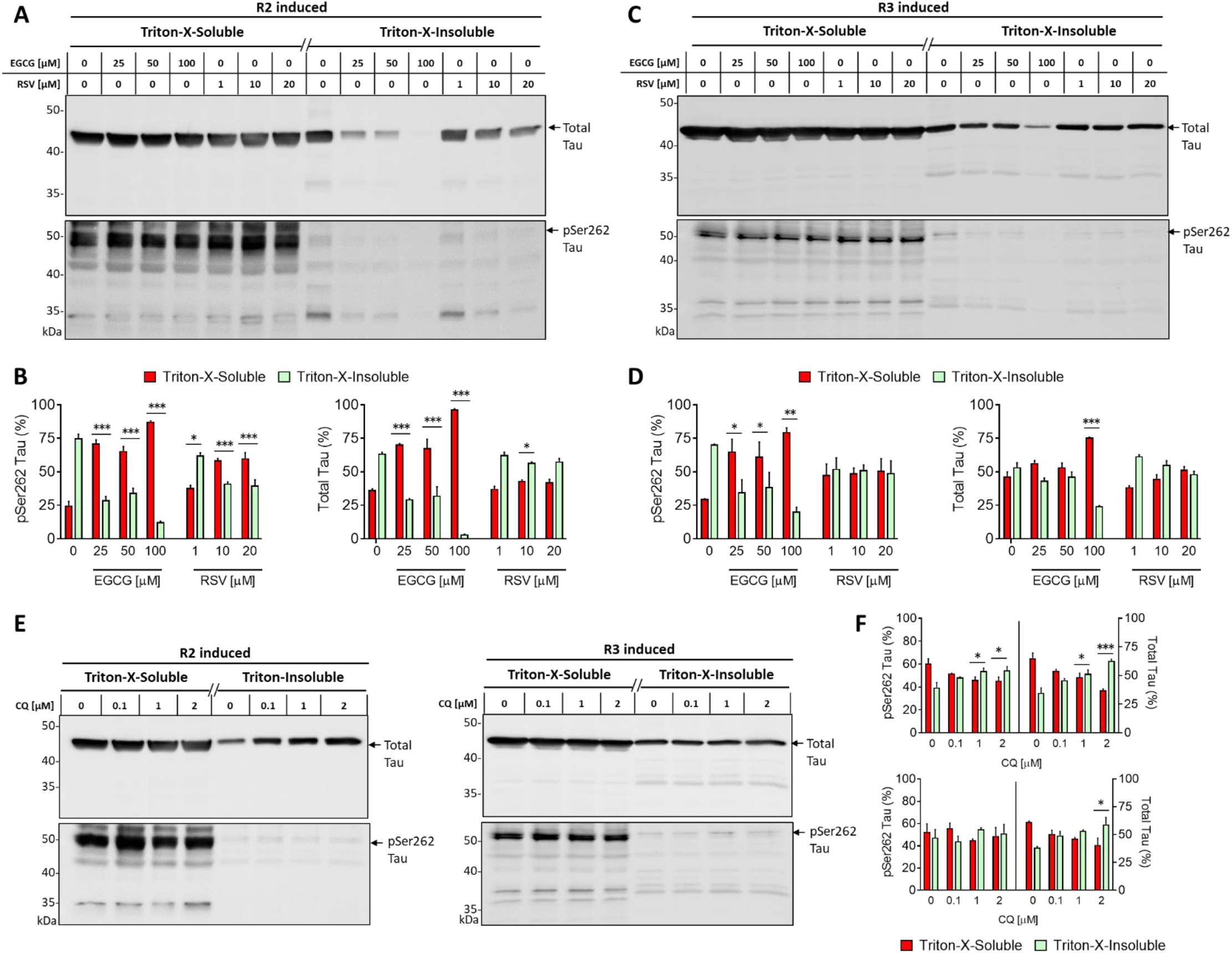
Autophagy promotes the removal of triton-insoluble total-tau and pSer262 tau aggregates. Western blots and quantitative analysis of triton-soluble and triton-insoluble fractions of pSer262 tau and total tau levels in cells induced with (A, B) R2 and (C, D) R3 fibrils after treatment with the indicated concentrations of EGCG and RSV for 48 hours or (E, F) in cells induced with R2 and R3 fibrils following treatment with CQ for 48 h. Data are presented as the mean ± SEM of 2 independent experiments. ***p < 0.001, **p < 0.01, *p < 0.05, Untreated (0 µM) vs Drug-treated, two-way ANOVA, Dunnet’s posthoc test. Images of uncropped blots are shown in the supplemental data (Fig S5).

Next, we used CQ to prove that autophagy failure results in proteotoxic stress and the buildup of intracellular aggregates of seeded tau. CQ significantly increased p62 levels in R2-induced cells and LAMP1 in R3-induced cells and inhibited the conversion of LC3-I to II in R2- and R3-induced cells (Fig. 4E, F). CQ-induced autophagy failure resulted in significant tau seeding under concomitant treatment conditions and showed a trend of increasing seeding when administered sequentially following cell induction with R3 fibrils (Fig. 3A).

The change in LAMP1 levels was identical in cells irrespective of whether cells were treated with autophagy inducer (EGCG) or blocker (CQ). In the case of EGCG, the increased LAMP1 levels correlate with the increased LC3A/B II/I ratio, corresponding to autophagy activation [36]. The increased LC3A/B II and LAMP1 levels were observed in the selective clearance of aberrant phosphorylated tau proteins by transcription factor EB [37]. This could prove the increased LAMP1 levels in EGCG-treated fibrils-induced cells. In the case of CQ-treated cells, the increased LAMP1 should correlate to the increased LC3A/B II/I ratio with defective autophagy due to the failure in the fusion of autophagosome and lysosome. Nevertheless, we observed a reduced LC3A/B II/I ratio and increased LAMP1 levels in CQ-treated cells. This still reflects a failure in autophagy induction, as similar results were obtained in the CQ-mediated cellular autophagy impairment [33].

### Enhancing autophagy clears triton-insoluble seeded pSer262 tau aggregates

Tau aggregates phosphorylated at Ser262 appear in the early AD phase [38]. We have recently shown that R2 and R3 cause a time- and dose-dependent increase in total- and pSer262- tau levels in triton-insoluble fractions of induced cells [28]. In addition, we have also shown that the presence of P301S mutation in biosensor cells enhanced pSer262 tau even in triton-soluble fractions, and P301S Tau is prone to aggregate into insoluble fractions upon exposure to exogenous R2 and R3 fibrils [27,28]. The pSer262-containing intracellular aggregates in R2 and R3-induced cells were also seed-competent [27]. Thus, the seed-competency of R2 and R3 fibrils is confirmed by the presence of insoluble factions of the total- and pSer262- tau in biosensor cells. Therefore, we next analysed triton-soluble and insoluble fractions of the total tau and pSer262 tau after EGCG, RSV and CQ treatment of the induced cells. EGCG significantly reduced triton-insoluble fractions of both total- and pSer262- tau in the induced cells, whereas RSV effects were noted only in R2-induced cells (Fig. 5A-D). In addition, EGCG was also able to inhibit the conversion of soluble pSer262-tau to insoluble pSer262-tau in the induced cells. Conversely, CQ treatment significantly increased total- and pSer262-tau levels of triton-insoluble fractions, albeit non-significantly, compared to non-induced cells (Fig. 5E, F). These results suggest that EGCG reduced triton-insoluble pSer262-tau levels in induced cells, possibly due to autophagy. In addition, EGCG may also impede the seed competency of fibrils, as it has been found to disaggregate AD brain-derived tau fibrils by binding to them [39]. These results further support our findings that autophagy enhancers effectively lower intracellular seeding by enhancing the autophagy-mediated clearance of pathologically phosphorylated (pSer262) aggregates of seeded tau.

## Discussion

Pathogenic tau aggregates promote intracellular tau seeding due to dysregulation or failure of innate cellular clearance systems such as the ALP [10]. Therefore, enhancing ALP could be an effective strategy to reduce the intracellular buildup of pathological aggregates and, thus, their spreading to naïve cells. Herein, we show that Tau R2 and R3 fibril-induced seeding produces proteotoxic stress, lowering autophagic activity, as evidenced by higher p62 levels and decreased conversion of LC3-I to -II, defining autophagy flux failure in fibril-induced biosensor cells.

Cells expressing P301L-tau have been shown to escape all three types of autophagy, such as macroautophagy, chaperone-mediated autophagy, and endosomal microautophagy-mediated degradation [18]. In addition, we have recently shown the enhanced conversion of soluble phosphorylated Ser262 tau into insoluble inclusions in R2 and R3-induced P301S-tau expressing cells [27,28]. In this study, we show that intracellular phosphorylated tau inclusions in R2- and R3-induced cells resist the aggregate-clearing mechanism and that failed autophagy is one of the mechanisms responsible for intracellular accumulation of aggregates. Interestingly, we also show a difference in the autophagic process between R2- and R3-induced cells, which might explain the pathological heterogeneity of tauopathies [40]. This difference in time-dependent autophagic failure and increased proteotoxic stress correlates with the difference in intracellular aggregate buildup between R2- and R3-induced cells, with R2- induced cells aggregating faster than R3-induced cells, as previously demonstrated [28]. Thus, our study shows that in addition to mutation, the seed competency of exogenous fibrils might also influence cellular clearance mechanisms, such as autophagy in removing intracellular tau aggregates in cells.

Autophagy modulators have been shown to clear the intracellular tau inclusions associated with neurodegenerative diseases [11,16,17]. Here, we show how exogenous R2 and R3 fibrils enhanced the already deteriorating autophagy process in cells expressing P301S tau and how autophagy induction rescued the cells from failed autophagic clearance of intracellular seeded tau. Furthermore, our study shows that EGCG-mediated clearance of intracellular tau aggregates results due to autophagy of insoluble phosphorylated tau inclusions. However, the effect of EGCG is more pronounced in R3-induced cells than in R2-induced cells in the early seeding phase. This indicates the difference between the seed-competency of R2 and R3 fibrils, which might affect the cellular clearance mechanisms against different pathogenic strains of tau.

Impairment of autophagy over time during ageing might contribute to the buildup of tangles of hyperphosphorylated tau [41]. We have observed that R3 fibril-seeded intracellular aggregates are more resistant to EGCG-mediated clearance than R2 fibril-induced intracellular aggregates at the later stage of seeding. Thus, there is the possibility that the increased buildup of tau tangles at the late stages of tauopathies might also be the contributing factor to the impairment in the already deteriorating autophagy process.

We found in our earlier study that intracellular seeding in cells induced with R3 fibrils corresponds to reduced cell growth [31]. In this study, we observed the decreased proliferation of R2 and R3-induced cells. A similar reduction in cell proliferation was noted in HEK293 cells containing aggregates of human FK506 binding protein 12 protein [42]. Differences among aggregate inheritance patterns were reported in various bacterial model systems. Protein aggregates are distributed equally between both daughter cells in *Caulobacter crescentus.* In contrast*, in E.coli* and mycobacteria, protein aggregates are retained in only one daughter cell type, and the other escapes aggregate inheritance [43]. A similar pattern of aggregate inheritance might play a pivotal role in eukaryotic cells in maintaining proteotoxic stress and its physiological consequences. Whether halting cell division is a defence mechanism for clearing or reducing the proteotoxic stress in aggregate-containing cells needs to be studied in detail.

In conclusion, our study explored tau seeding using Tau R2 and R3 fibrils, revealing that differences in seeding dynamics are tied to autophagy failure. This could explain the varied pathology and progression between 3R and 4R tauopathies, especially in 3R tau isoforms without R2. Enhancing autophagy with EGCG cleared intracellular tau aggregates and reduced pSer262 tau aggregates, offering potential treatments for specific tauopathies, particularly those without R2 in 3R tau isoforms.

## Materials and methods

### Cell culture, chemicals and reagents

HEK293T Tau RD P301S FRET biosensor cells, hereafter referred to as only biosensor cells for simplicity, were purchased from the American Type Culture Collection (Tau biosensor cells; ATCC® CRL-3275™) and cultivated in complete Dulbecco’s Modified Eagle’s growth medium (Lonza, Cat. # 12–604F) supplemented with 10% fetal bovine serum (Gibco, Cat. # 10270), 1% antibiotics (streptomycin and penicillin) (Diagnovum, Cat. # 910) and 1% GlutaMax (Gibco, Cat. #35050-061) at 37 °C in a humidified 5% CO_2_ incubator. The cell line was regularly tested for bacterial, fungal and mycoplasma contaminations and authenticated biweekly or monthly.

Autophagy modulators: Epigallocatechin gallate (Cat. # E4143), resveratrol (Cat. # R5010), L-Ascorbic acid (Cat. # A4403), quercetin (Cat. # Q495), carbamazepine (Cat. # C4024), artemisinin (Cat. # 361593); dexamethasone (Cat. # D4902), chloroquine (Cat. # C6628) and other chemicals, unless otherwise mentioned, were purchased from Sigma-Aldrich (St. Louis, MO, USA).

### Tau seeding assay for protein extraction

Biosensor cells were plated in a 6-well plate (0.5 x 10^6^ cells/mL, 2 mL of medium /well) and cultured at 37 °C in a 5% CO_2_ incubator overnight. R2 and R3 fibrils were prepared by aggregating for 48 hours [27]. The next day, cells were transfected with 50 nM fibrils collected from the *in vitro* aggregation assay, carried out without Thioflavin T, using Lipofectamine™ 3000 Transfection Reagent. First, 50 nM of fibrils were mixed in Opti-MEM™ I Reduced Serum Medium (Gibco, Cat. # 11058021) supplemented with P3000™ reagent (1.5 µL/well) and Lipofectamine™ 3000 reagent (3 µL/well). The transfection mixture was incubated for 15 minutes at room temperature before adding to cells. Next, the old growth media was replaced with pre-warmed fresh complete growth media, and the transfection mixture was added dropwise into wells while swirling the content of the plate. After 6 hours, the transfection media was removed, and the cells were replaced with pre-warmed fresh complete media. Cells were collected at 3, 6, 9, 12, 24, 36, and 48 hours post-transfection and lysed as described below. The cells were concomitantly treated with 50 nM fibrils and selected concentrations of compounds for 48 hours for compound treatment.

When collecting cells, the old medium was collected in a 15 mL Falcon tube to collect any detached cells, centrifuged at 300 x g for 5 minutes to pellet, and washed with 1× Tris-buffered saline (TBS, pH 7.4). Next, the remaining attached cells were collected following cell detachment using 200 µL of TrypLE (Gibco) and washing with a complete growth medium. Next, the cell suspension was collected in the same 15 mL Falcon tube containing detached cells and centrifuged at 300 x g for 5 minutes. The overall effect of fibrils on cellular seeding and autophagy was analysed by collecting cells, including detached and attached populations. Next, the complete cell pellet was resuspended in 1 mL of 1 x TBS, transferred into a 1.5 mL centrifuge tube, and re-centrifuged for 10 minutes at 500 x g. Finally, the supernatant was discarded, and the sedimented pellet was processed for protein extraction, as described below.

### Sorting of R2 and R3 fibril-induced cells

Briefly, cells were collected in a growth medium, and the cell density was adjusted to a final concentration of 10 x 10^6^ cells/mL. Cells were analysed on FACSAria II (Becton Dickinson) cell sorter. First, 5 x 10^4^ cells were acquired to set up gating. Dead cells and debris were gated out by forward scatter versus side scatter gating (Fig. 2A). Then, intact cells were gated and brought to the blue/green channels dot-plot diagram. The gating strategy used to sort R2/R3 fibril-induced seeded cells into different fractions containing 0 % or 100 % of Ag+ cells was based on the different expression intensities of particular fluorescent protein tags corresponding to plasmids used for transfection (Fig. S1). Green fluorescent marker was excited by a 488 nm laser line and the fluorescence signal was collected at 525 nm (green) channel. The blue fluorescent marker was excited by a 445 nm laser line, and the fluorescence signal was collected with a 530 nm channel. Fractions containing 0 % or 100 % of Ag+ cells were collected in FACS tubes at 4°C by two-way sorting mode setting. The tubes were immediately put on ice and proceeded for western blot.

### Total protein extraction and triton X-100 cell fractionation

Total proteins were extracted by cell lysis in RIPA lysis buffer (Thermo Scientific, Cat. # 89901) supplemented with protease (Roche, Cat. # 04693116001) and phosphatase (Roche, Cat. # 04906837001) inhibitors. Briefly, the cell pellet was suspended in 100 µL lysis buffer and mixed gently, followed by sonication in a water bath sonicator for 3 minutes at 25% Amplitude with a 15-sec pulse ON and 15-sec pulse OFF. The samples were next centrifuged at 12 000 RPM at 4 °C for 20 minutes, and the supernatant was collected in pre-chilled 1.5 mL fresh Eppendorf tubes and labelled as total protein extracts. The remaining cell pellet was discarded.

Triton X-100 cell fractionations were prepared as described previously [27]. Briefly, cells were harvested and resuspended in 1× TBS containing 0.05% triton X-100 supplemented with protease and phosphatase inhibitors. The cell lysate was clarified by centrifuging at 500 × g for 5 minutes, followed by re-centrifuging at 1000 × g for another 5 minutes at 4 °C on a benchtop centrifuge. Next, the supernatant was collected and centrifuged at 65,000 × g for 30 minutes at 4 °C to collect the supernatant as the soluble fraction. The pellet was resuspended in RIPA buffer supplemented with protease and phosphatase inhibitors and collected as the insoluble fraction. Protein concentration in samples was quantified using a BCA protein assay kit, following the manufacturer’s instructions (ThermoFisher Scientific) before processing by immunoblotting, as described below.

### SDS-PAGE electrophoresis and Western blotting

Thirty-five µg of total protein and 45 μg of triton X-100 soluble fraction and an equal volume of insoluble fractions were electrophoresed by 10% or 12% SDS-PAGE at 200 Volts. Electrophoresed proteins were transferred onto a 0.2 µm nitrocellulose membrane (Bio-Rad, Hercules, CA, United States) using a Trans-Blot Turbo Transfer System (Bio-Rad) for 10-15 minutes at 20-25 Volts. Transferred membranes were blocked in 5% BSA in 1x TBS-Tween- 20 (TBST) buffer for 1 hour and incubated with primary antibodies overnight at 4 °C. Protein bands were developed using Alexa 488 fluorescent dye-conjugated secondary antibodies for 2 hours at room temperature. Images of membranes were acquired on a Bio-Rad Gel Doc XR + Gel Documentation System, and processed using ImageJ (NIH) software, as described elsewhere [27]. Primary antibodies used were anti-LC3A/B (rabbit polyclonal; 1: 1000 dilution; Invitrogen, Cat. # PA1-16931), anti-p62 (mouse monoclonal; 1: 1000 dilution; CST, Cat. # 88588), anti-LAMP1 (rabbit monoclonal; 1: 1000 dilution; CST, Cat. # 9091), anti-total tau (mouse monoclonal; 1: 1000 dilution; Sigma-Aldrich, Cat. # 05-804), anti-phospho Ser262 tau (rabbit polyclonal; 1: 1000 dilution; Invitrogen, Cat. # PA5-85654), and anti-β actin (mouse monoclonal; 1:4000 dilution; Sigma-Aldrich, Cat. #A2228). Secondary antibodies used donkey anti-mouse IgG (H+L) Alexa Fluor™ 488 (1:2000; Cat. #A21202, Invitrogen) and goat anti-rabbit IgG (H+L) Alexa Fluor™ 488 (1:2000; Cat. #A11034, Invitrogen).

### Drug treatment and tau seeding assay

Biosensor cells were plated at a density of 0.15 x 10^6^ cells/mL in a CellCarrier 96-well plate (PerkinElmer, Cat. # 6005550). Cells were transfected with 50 nM R2/R3 fibrils premixed in Opti-MEM™ I Reduced Serum Medium supplemented with P3000™ reagent (0.25 µL/well) and Lipofectamine™ 3000 reagent (0.5 µL/well). Cells were treated with drugs by two different methods: (1) Sequential treatment: the cells were first transfected with 50 nM R2/R3 fibrils for 6 hours, rinsed with 1x PBS to remove residual transfectants and then treated with the drugs at different concentrations for 12 hours, and (2) Concomitant treatment: the cells were treated with 50 nM R2/R3 fibrils in the presence or absence of compounds at different concentrations for 24 hours. The seeding of native tau in cells was measured by imaging, as described below.

To find the effect of compounds (autophagy inhibitor (CQ)/inducer (EGCG), proteasome inhibitor (BTZ, Bortezomib) and protein synthesis inhibitor (CHX, Cycloheximide), which reported to suppress autophagy-mediated protein degradation on the clearance of intracellular tau aggregates post-seeding, the biosensor cells were plated at a density of 0.075 x 10^6^ cells/mL in a CellCarrier 96-well plate (PerkinElmer, Cat. # 6005550). As described above, cells were transfected with 50 nM R2/R3 fibrils. After 48 hours of transfection, the transfection mixture containing fibrils was removed and washed with 1x PBS, and then the seeded cells were treated with CQ (1, 2 µM), EGCG (50, 100 µM), BTZ (5, 10 nM) and CHX (100, 200 nM) for 72 hours. The imaging of the level of intracellular tau aggregates was measured as mentioned below.

### Imaging and quantification of tau seeding

The old growth media was replaced with phenol red-free growth media, and the cells were imaged on a Cell Voyager CV7000S microscope (Yokogawa, Tokyo, Japan) using a 20x objective, as described previously [27]. Before imaging, biosensor cells were stained with 10 μM Hoechst 33342 nuclear dye (Invitrogen, Cat. #H21492) for 10–15 min at 37 °C. The intracellular CFP/YFP Tau RD P301S aggregates and cell numbers were quantified using Columbus Image Data Storage and Analysis System (PerkinElmer), as described before [27].

### Cytotoxicity assay

Biosensor cells were plated at a density of 1 x 10^5^ cells/mL in 96-well plates and incubated overnight at 37 °C in a 5% CO_2_ incubator. The next day, cells were treated with drugs for 48 hours, and the viability was assessed by a standard MTT assay. The half-maximal inhibitory concentrations (IC_50_) of compounds were calculated using GraphPad Prism Software (Trail Version; San Diego, CA, USA).

### Statistical analysis

All experiments were repeated at least 2 independent times. All statistical analyses were performed in Statistica Version 14 (TIBCO Software Inc., CA, USA), and differences were considered significant at P < 0.05.

## CRediT authorship contribution statement

**Narendran Annadurai:** Conceptualisation, Investigation, Methodology, Writing – original draft; **Agáta Kubíčková:** Methology; **Ivo Frydrych:** Methology; **Marián Hajdúch**: Funding acquisition, Supervision, Writing; **Viswanath Das:** Conceptualisation, Investigation, Funding acquisition, Writing - Review & Editing, Supervision.

## Acknowledgments

This work was supported in parts by the Internal Student Grant Agency of the Palacký University in Olomouc, Czech Republic (IGA_LF_2022_033), infrastructural projects (CZ-OPENSCREEN – LM2023052; EATRIS-CZ – LM2023053), the projects National Institute for Cancer Research (Program EXCELES, ID Project No. LX22NPO5102) and National Institute for Neurological Research (Program EXCELES, ID Project No. LX22NPO5107) - Funded by the European Union - Next Generation EU from the Ministry of Education, Youth and Sports of the Czech Republic (MEYS), and the Grant Agency of the Czech Republic (23- 06301J).

## Data statement

The raw data supporting this study’s findings are available upon reasonable request from the corresponding author (V Das).

## Abbreviations

AD: Alzheimer’s disease
ALP: autophagy-lysosome pathway
BTZ: bortezomib
CQ: chloroquine
CHX: cycloheximide
EGCG: epigallocatechin gallate
LAA: l-ascorbic acid
R2: repeat 2
R3: repeat 3
RSV: resveratrol
QCT: quercetin.

