## Supplementary Figure S1-S5 for "Differential seeding by exogenous R2 and R3 fibrils influences autophagic degradation of intracellular tau aggregates in Tau K18 P301S cells"

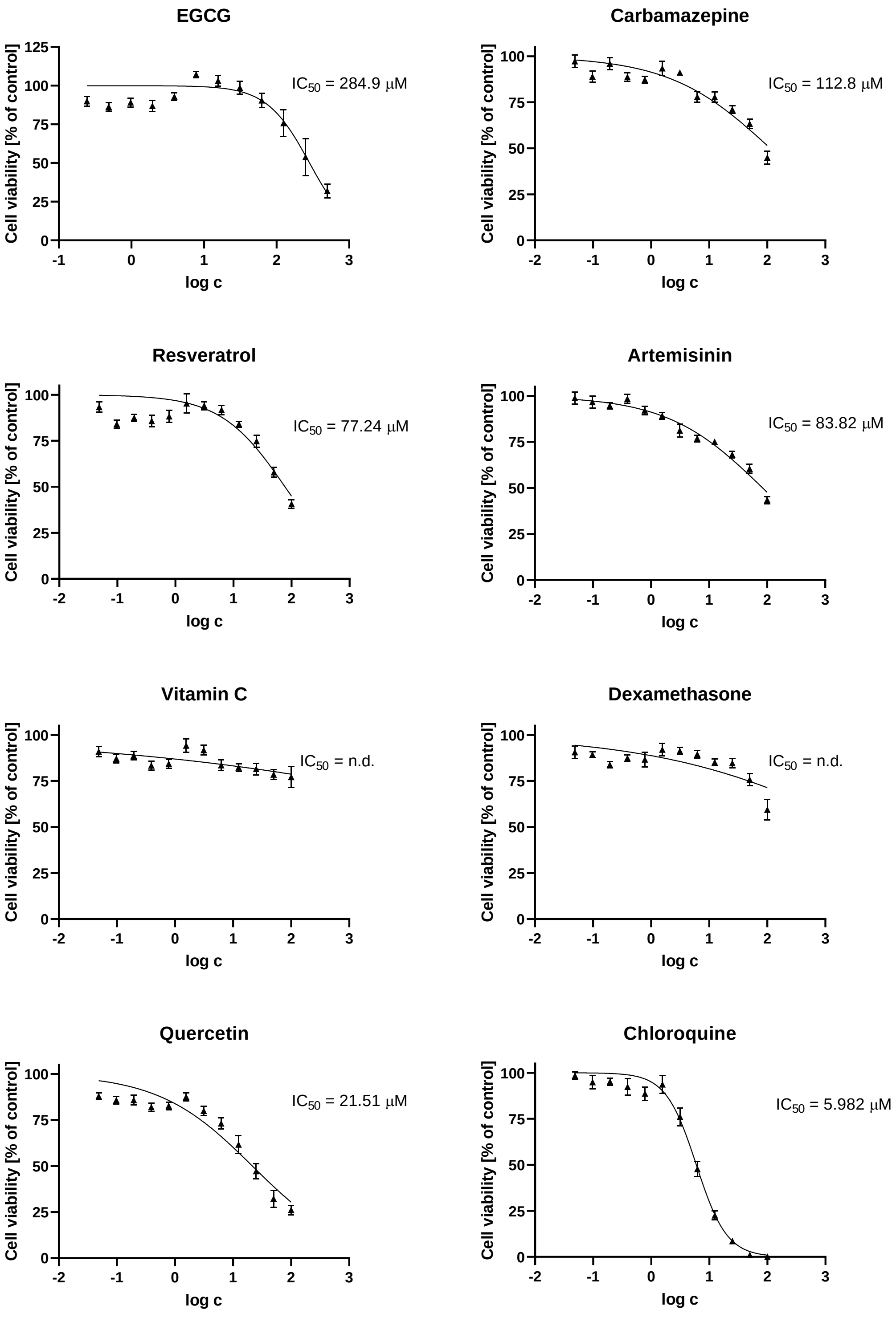


**Figure S1.** Dose-response curves of compounds and the calculated $IC_{50}$ concentrations are shown. N.d., Not determined. Data are presented as the mean ± SEM of 2 independent experiments.

**Uncropped images of blots shown in Figure 1A (R2 induced)**

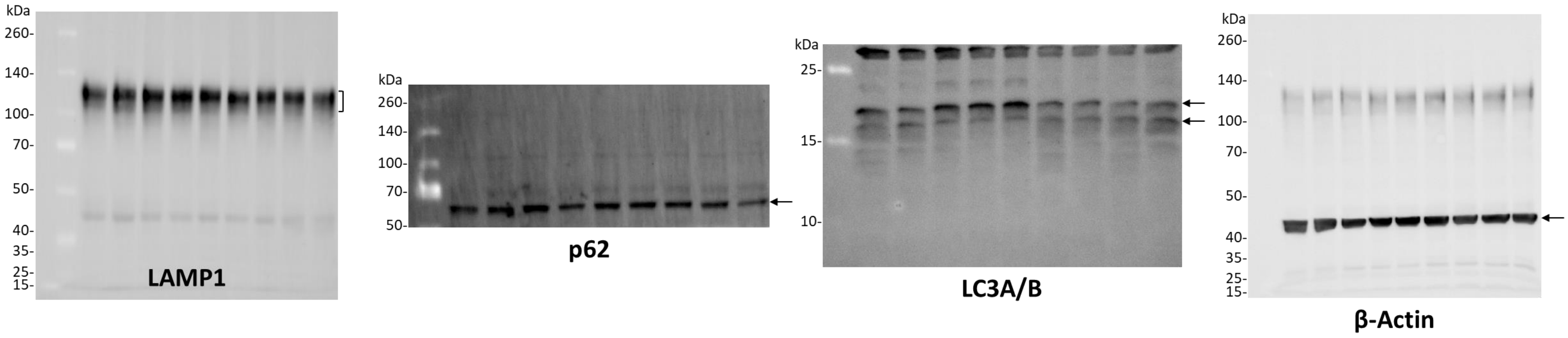


**Uncropped images of blots shown in Figure 1B (R3 induced)**

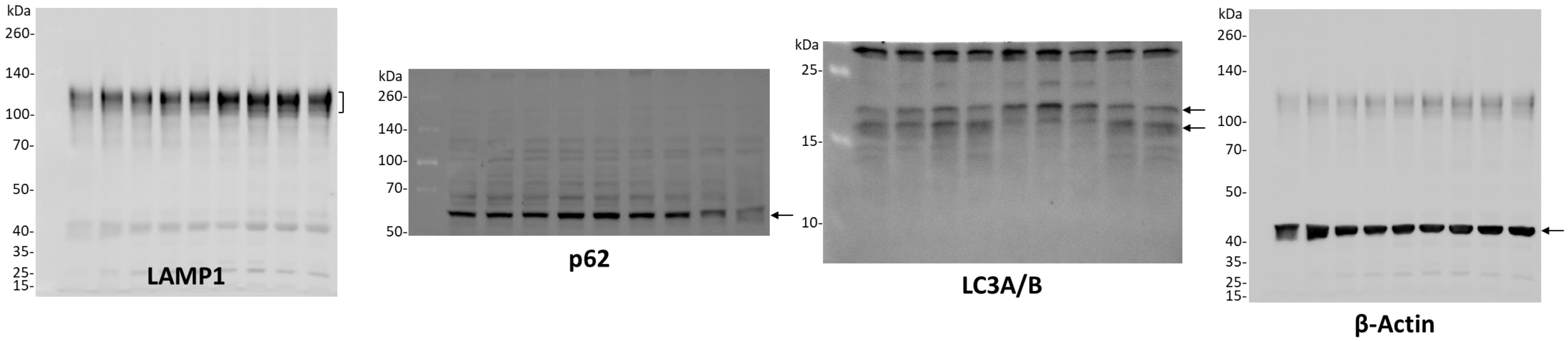


**Figure S2**. Images of uncropped blots that were presented in Figure 1 are shown here. Note that for p62 and LC3A/B, blots were horizontally cut, and each separate blot was stained with different primary antibodies.

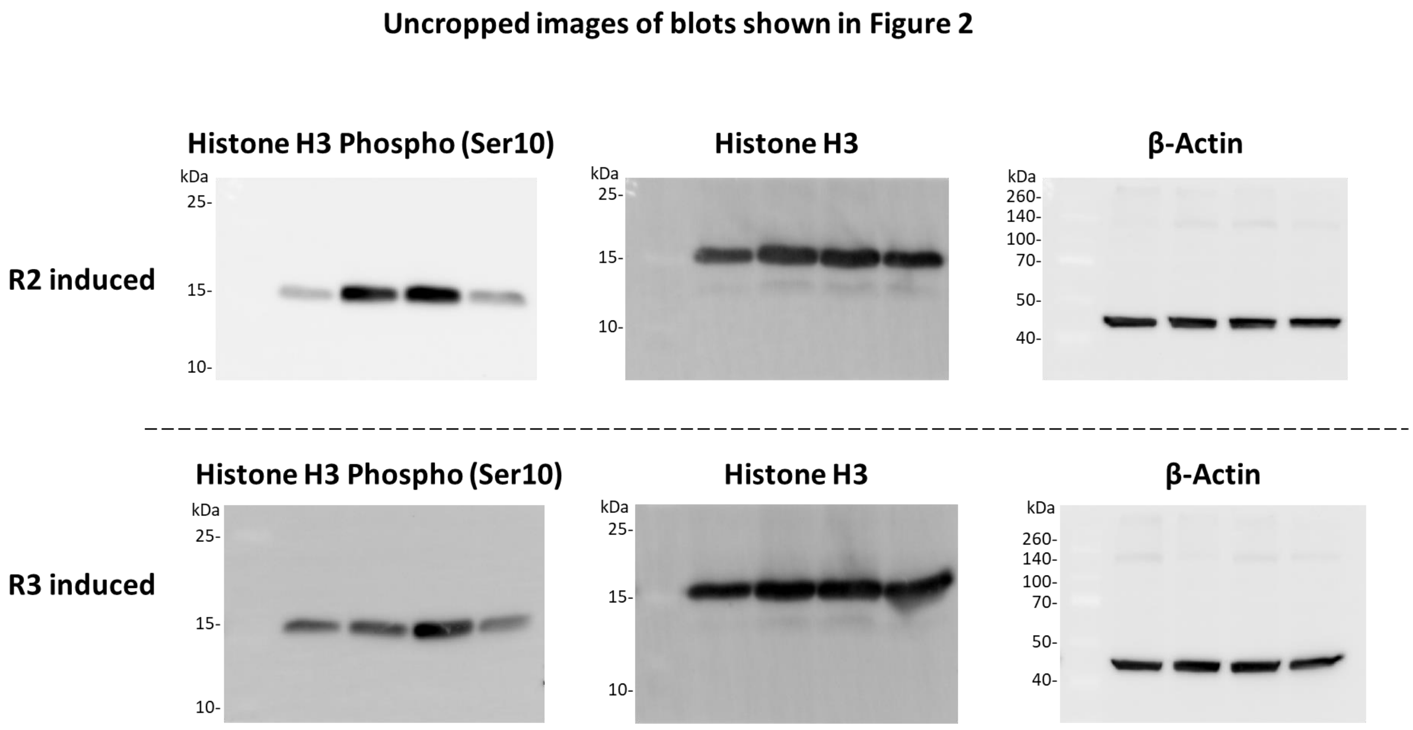


**Figure S3**. Images of uncropped blots that were presented in Figure 2 are shown here. Note that blots were horizontally cut, and each separate blot was stained with different primary antibodies.

**Uncropped images of blots shown in Figure 4**

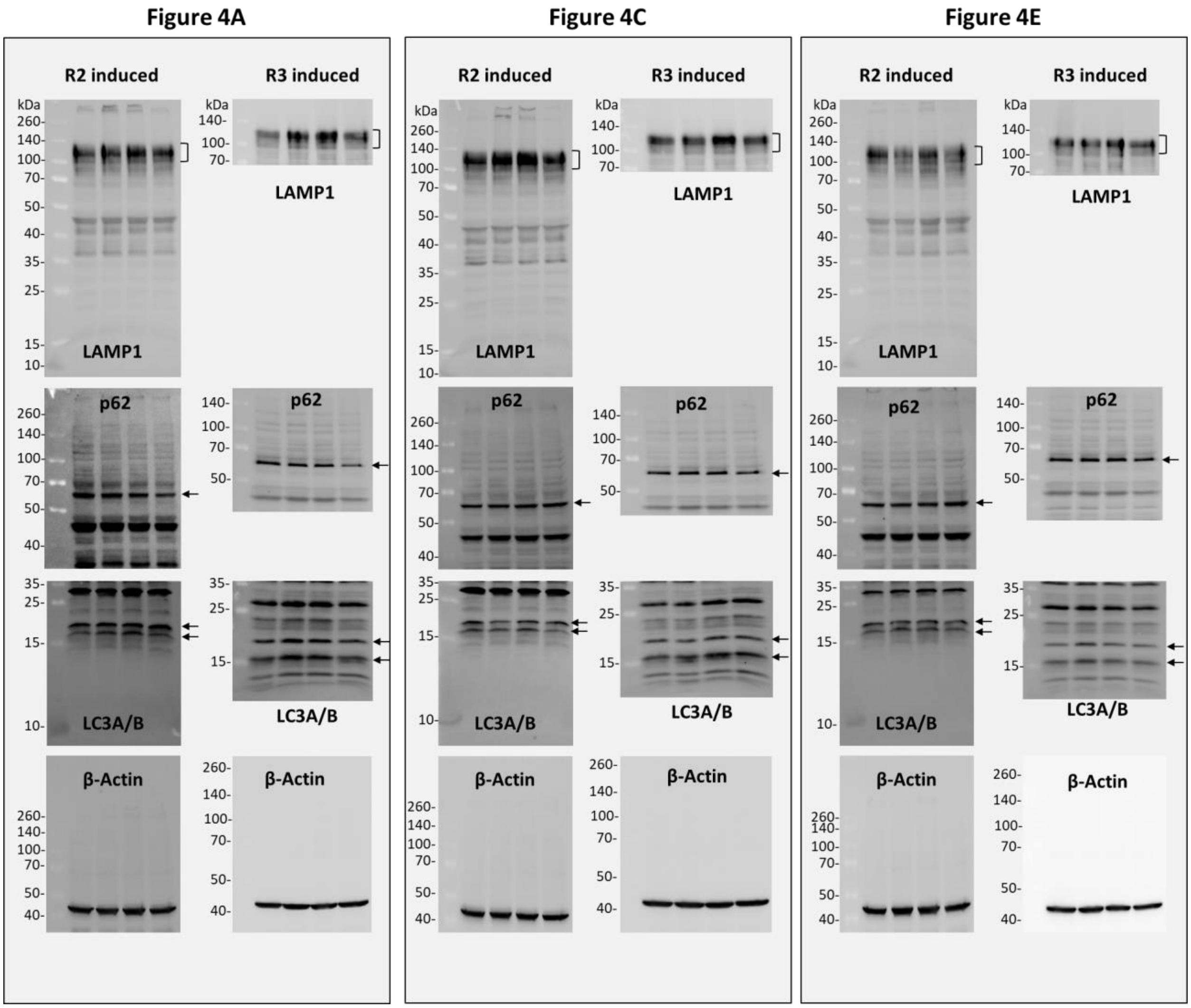


**Figure S4**. Images of uncropped blots that were presented in Figure 4 are shown here. Note that blots were horizontally cut for some experiments, and each separate blot was stained with different primary antibodies.

### Uncropped images of blots shown in Figure 5

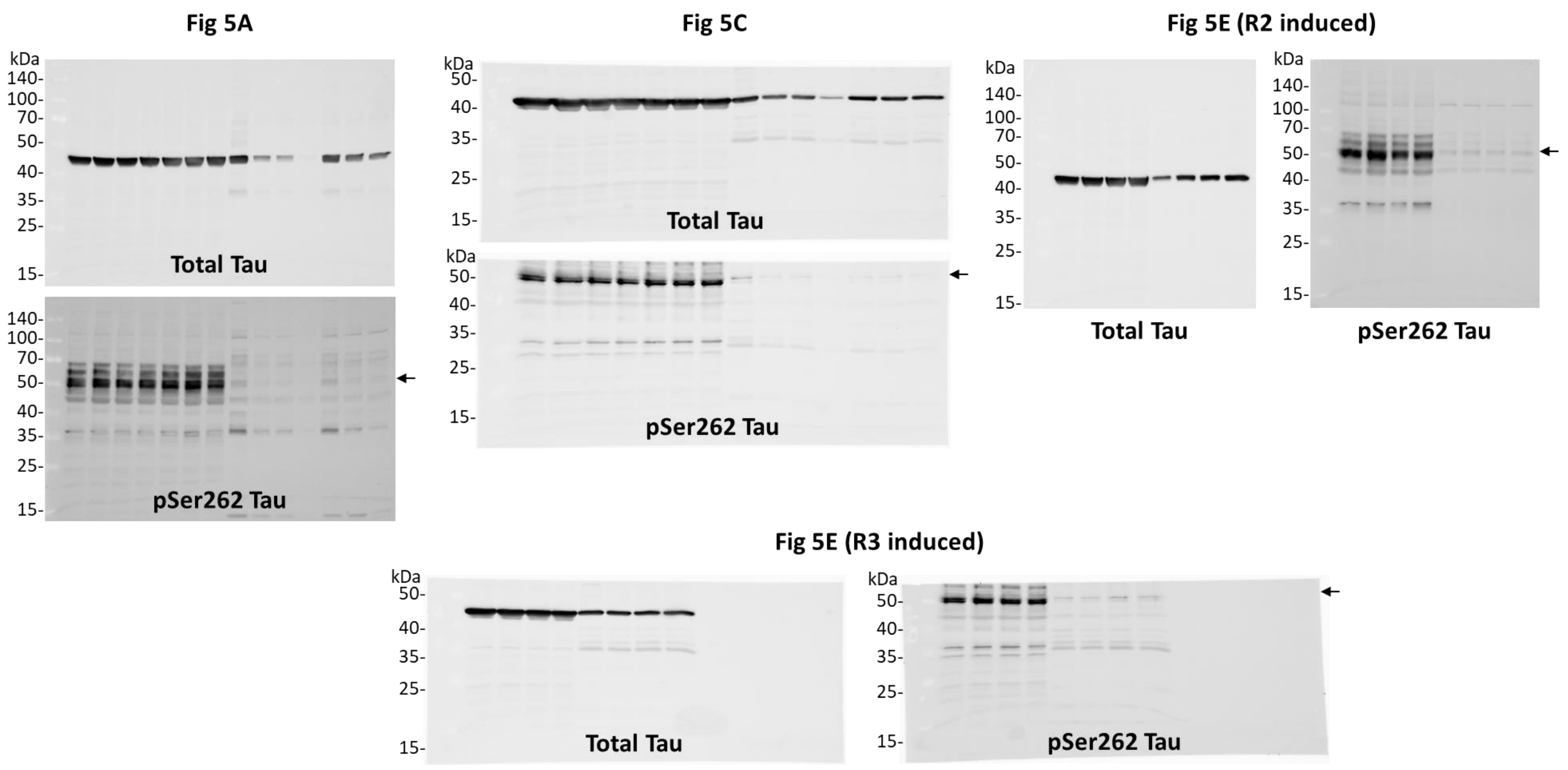


**Figure S5**. Images of uncropped blots that were presented in Figure 5 are shown here. Note that blots were horizontally cut for some experiments [Fig 5C and 5E (R3 induced)], and each separate blot was stained with different primary antibodies.
